# Adaptive thermogenesis reprograms the behavioural and metabolic responses to protein restriction for successful amino acid homeostasis

**DOI:** 10.1101/2025.07.30.667623

**Authors:** Anthony H. Tsang, Simon Benoit, Sam Lockhart, Ana Parish, Sam Virtue, Lorena da Silva Paes, Xue Li Wong, Debra Remmington, Antonio Vidal-Puig, Stephen O’Rahilly, Anthony P. Coll, Valentina Pirro, Albert Koulman, Clemence Blouet

## Abstract

Reduced protein intake is proposed to contribute to the obesity epidemic, but existing murine models of protein restriction do not promote obesity, limiting insight into underlying mechanisms. Here we show that mice self-select a consistent daily protein intake, independent of energy needs. Under mild protein restriction (7-10% protein), mice exhibit hyperphagia and increased adiposity. This hyperphagic response is blunted at thermoneutrality, leading to loss of lean mass and body weight. Protein-restricted mice also fail to exhibit protein preference at thermoneutrality, despite elevated circulating FGF21 levels. Circulating levels of essential amino acids (AAs) are tightly regulated during protein restriction at 22°C, but this regulation is lost at 28°C. Metabolomic and transcriptional analyses revealed a role for hepatic and brown fat AA-derived acylcarnitines and N-acetyl AAs in buffering free AA levels for the maintenance of AA homeostasis at 22°C, but these pathways are blunted at thermoneutrality. Thus, the behavioural and metabolic adaptations to protein restriction rely on coordinated peripheral and central mechanisms, modulated by ambient temperature.

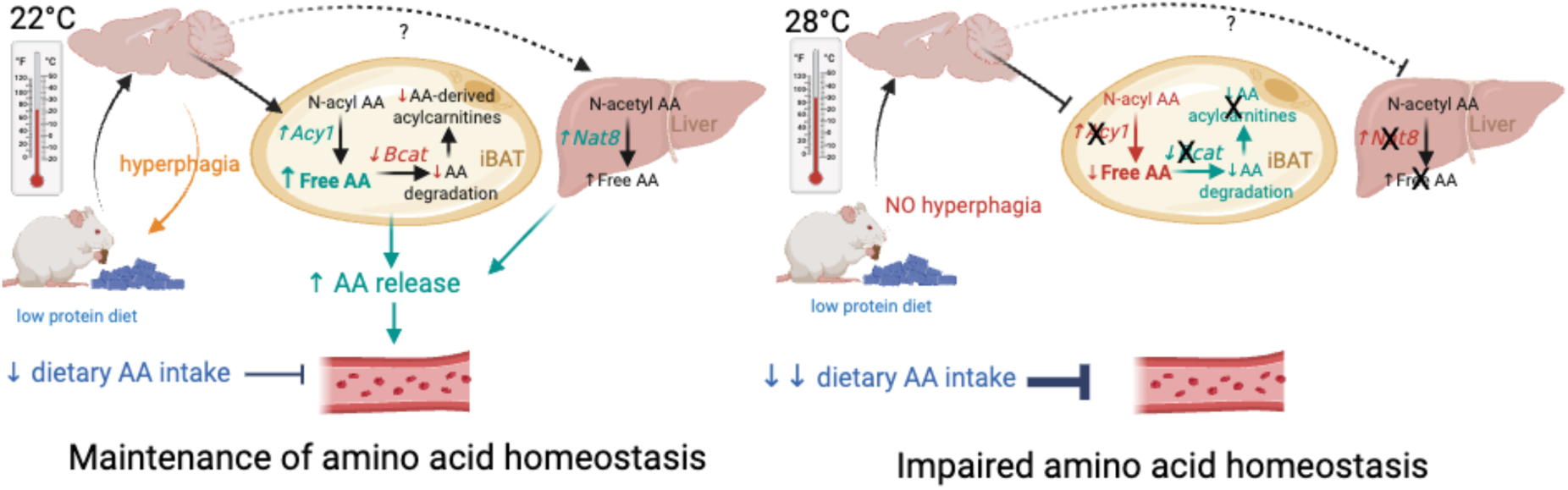

## INTRODUCTION

Half of the amino acids (AAs) needed for physiological functions cannot be synthesized by mammals and must be obtained through the diet, but how organisms adapt to suboptimal dietary protein intake to maintain systemic AA availability remains unclear. In a variety of experimental models, dietary protein restriction associates with hyperphagia ^1,2 3^. Consistently, healthy humans report increased hunger and energy intake during exposure to low-protein diets ^4,5^. This observation is the foundation of the protein leverage hypothesis, which predicts that animals eat until they consume enough protein to meet bodily needs, leading to overeating when dietary protein content is too low ^1,2, 6^. Protein leverage is proposed as a mechanism contributing to the obesity epidemic ^7–10^, but evidence for causality is lacking.

Progress on this question has been hindered by the lack of experimental model of obesity induced by dietary protein restriction. In fact, mice fed a low-protein diet exhibit weight loss despite significant hyperphagia ^11–13^. This phenotype is observed following a restriction of dietary protein intake to 5% or less of daily energy intake, which associates with reduced lean mass and hypoalbuminemia, suggesting protein malnutrition ^14–16^. The feeding and metabolic responses observed in this context require brain FGF21 signalling ^11^ and primarily reflect the hypermetabolic response to elevated FGF21 levels. However, it is unclear whether alternative pathways might be recruited in response to milder protein restriction, allowing the maintenance of systemic AA availability, lean mass and normal growth.

In this study, we designed experiments to test the control of dietary protein intake in male mice and characterise the feeding and metabolic responses to mild protein restriction. We housed mice at 22°C (standard laboratory conditions) and thermoneutrality (28°C) to clarify the role of thermoregulatory energy expenditure in the metabolic response to protein restriction. We characterised the consequences of mild protein restriction on systemic AA availability and identified mechanisms recruited to maintain circulating AA levels during dietary protein restriction.

## Results

### Protein intake is controlled independently of energy intake in mice

The protein leverage hypothesis is built on the premise that dietary protein intake is controlled to reach a daily protein intake target, leading to hyperphagia and weight gain when dietary protein availability is low ^6^. We first asked whether under standard laboratory conditions, young adult C57/Bl6J males maintain dietary protein intake to a constant level. Mice fed a normal chow diet were exposed to two isocaloric, isofat diets offered ad libitum for 5 days and containing either 5% protein and 75% carbohydrate (low protein, P5), or 45% protein and 35% carbohydrate (high protein, P45) (**Fig. 1A**). Under these conditions, energy intake is dissociated from carbohydrate and protein intake, and mice can self-select the carbohydrate: protein ratio of their diet. Although daily energy intake varied between mice (9 to 16 kcal), as did carbohydrate and fat intake, mice ate on average 1.9±0.04 kcal protein/day, suggesting a tight control of protein intake (**Fig. 1B**).

**FIGURE 1:**
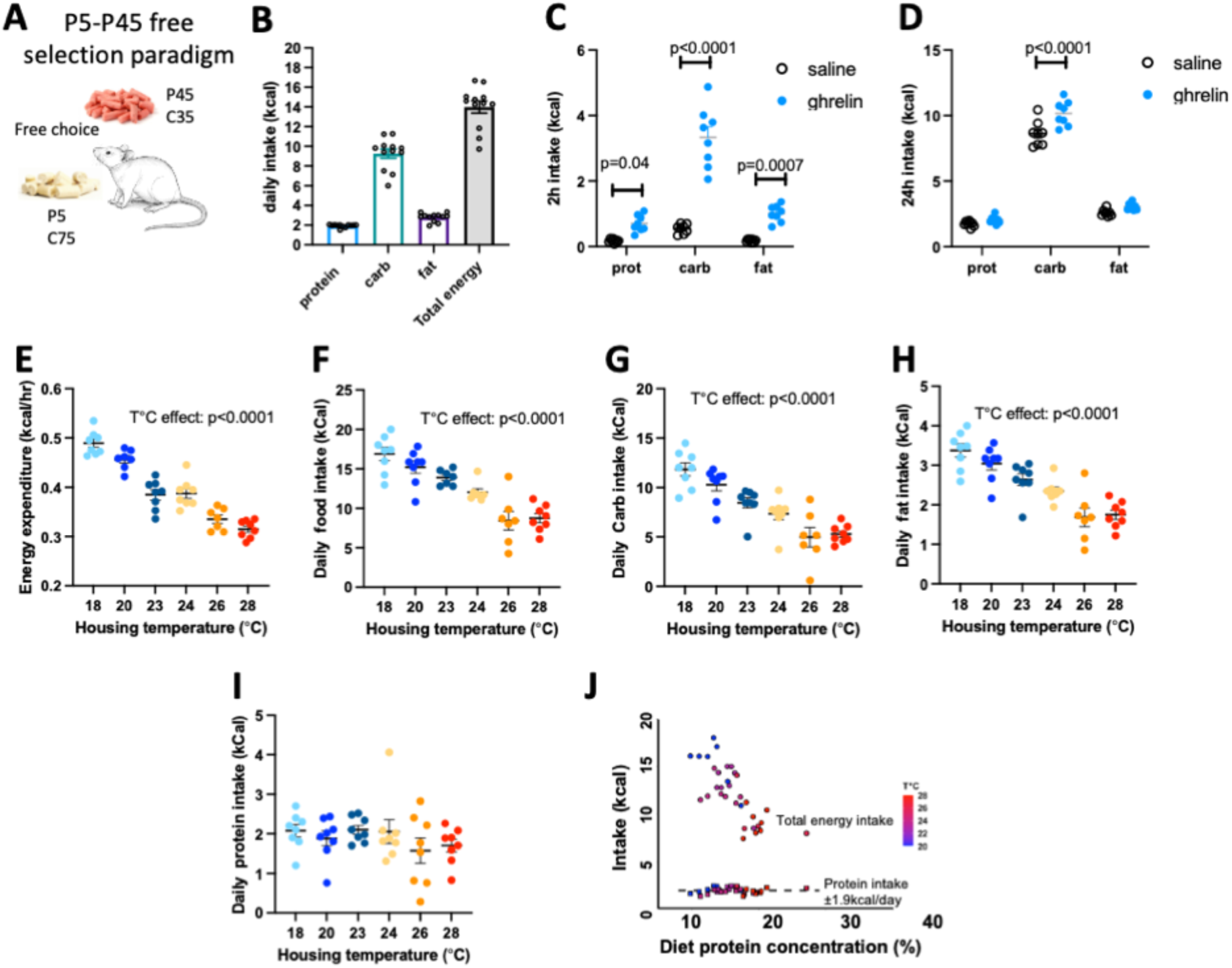
Protein intake is regulated in mice independently of energy intake. (**A**)The P5-P45 free selection paradigm: mice have ad libitum access to the P5 (5% protein, 75% carbohydrate) and the P45 (45% protein, 35% carbohydrate) diets. (**B**) Macronutrient and energy intake in male mice housed at 24°C and fed ad libitum (n=8). (**C, D**) Macronutrient intake 2h (**C**) and 24h (**D**) following an ip injection of vehicle or ghrelin in mice exposed to a P5-P45 free selection paradigm (n=8). (**E-J**) Energy expenditure (**E**), energy intake (**F)**, carbohydrate intake **(G),** fat intake (**H)** and protein intake (**I**) in mice exposed to a P5-P45 free selection paradigm and housed in a temperature gradient (n=8). (**J**) Correlations between energy/protein intake and dietary protein concentration at various housing temperatures. Data (means± sem) were analysed using a 2-way ANOVA for repeated measures.

If protein intake is controlled to meet the body’s protein needs independently of energy intake, then protein intake should remain stable under conditions of increased or decreased energetic demand. To test this, we exposed mice to the P5-P45 free selection paradigm as above (**Fig. 1A**) following an intraperitoneal (ip) injection of saline or the orexigenic hormone ghrelin. Consistent with the hunger-promoting properties of ghrelin, ghrelin induced a significant increase in energy intake primarily driven by an increase in carbohydrate intake, while protein intake remained similar between vehicle- and ghrelin-treated groups, reaching an average of 1.9 kcal after 24h (**Fig. 1C, 1D**).

Next, we took advantage of the fact that murine energetic needs are strongly and linearly correlated with housing temperature. We housed mice in a thermal gradient (28°C to 18°C) to produce a wide range of energetic needs. As expected, energy intake and energy expenditure increased linearly when decreasing housing temperature (**Fig. 1E, 1F**). Changes in energy intake were driven by changes in fat and carbohydrate intake, while protein intake remained constant (**Fig. 1F-1I**).

Consequently, at lower housing temperature, the concentration of protein in the diet decreased while total energy intake increased (**Fig. 1J**). Body weights remained steady across the temperature gradient, indicating successful maintenance of energy balance under these conditions (not shown). Collectively, these data suggest that when protein and energy intake can be regulated independently, protein intake is maintained constant, reaching an average daily value of 1.9 kcal in young adult C57/bl6J mice. Accordingly, we conclude that in standard laboratory housing conditions at 22-24°C, the optimal protein concentration is 14%. However, because most animal houses routinely use diets containing 20% protein, we used 20% protein in control diets in all subsequent experiments.

### Mild protein restriction increases energy intake and adiposity but does not increase weight gain in male mice

In the P5-P45 free selection paradigm, protein and energy intake are controlled independently, and both energy and protein needs can be met without trade-off, but that’s rarely the case in the wild. We next focused on testing how energy and protein needs are prioritised in conditions where protein availability is suboptimal. In the literature, this question has often been investigated using diets containing 5% protein. Under these conditions, mice lost lean mass despite sustained hyperphagia, suggesting that under these conditions, dietary protein intake is insufficient, compromising systemic AA homeostasis^9^. Thus instead, we tested the effect of mild protein dilution (10% protein diet – P10).

Mice maintained on the P10 diet consistently increased their daily energy intake (+32% on average) compared to controls fed a diet containing 20% protein (P20) (**Fig. 2A**). Despite this hyperphagia, protein intake remained significantly lower in P10-fed than in P20-fed mice (**Fig. 2B**), but this deficit did not prevent weight gain (**Fig. 2C**) and was not associated with reduced lean mass (**Fig. 2D**). Mice on P10 gained significantly more fat mass over 28 days (**Fig. 2E**). Energy expenditure did not significantly differ between groups (**Fig. 2F, 2G**), despite elevated circulating FGF21 levels (**Fig. 2H**). We measured a strong trend towards elevated day-time respiratory quotient (RQ) in P10-fed mice, indicating altered substrate utilisation (**Fig. 2I, 2J**). Thus, mild protein restriction produces hyperphagia and increases adiposity but does not lead to significant weight gain.

**Figure 2:**
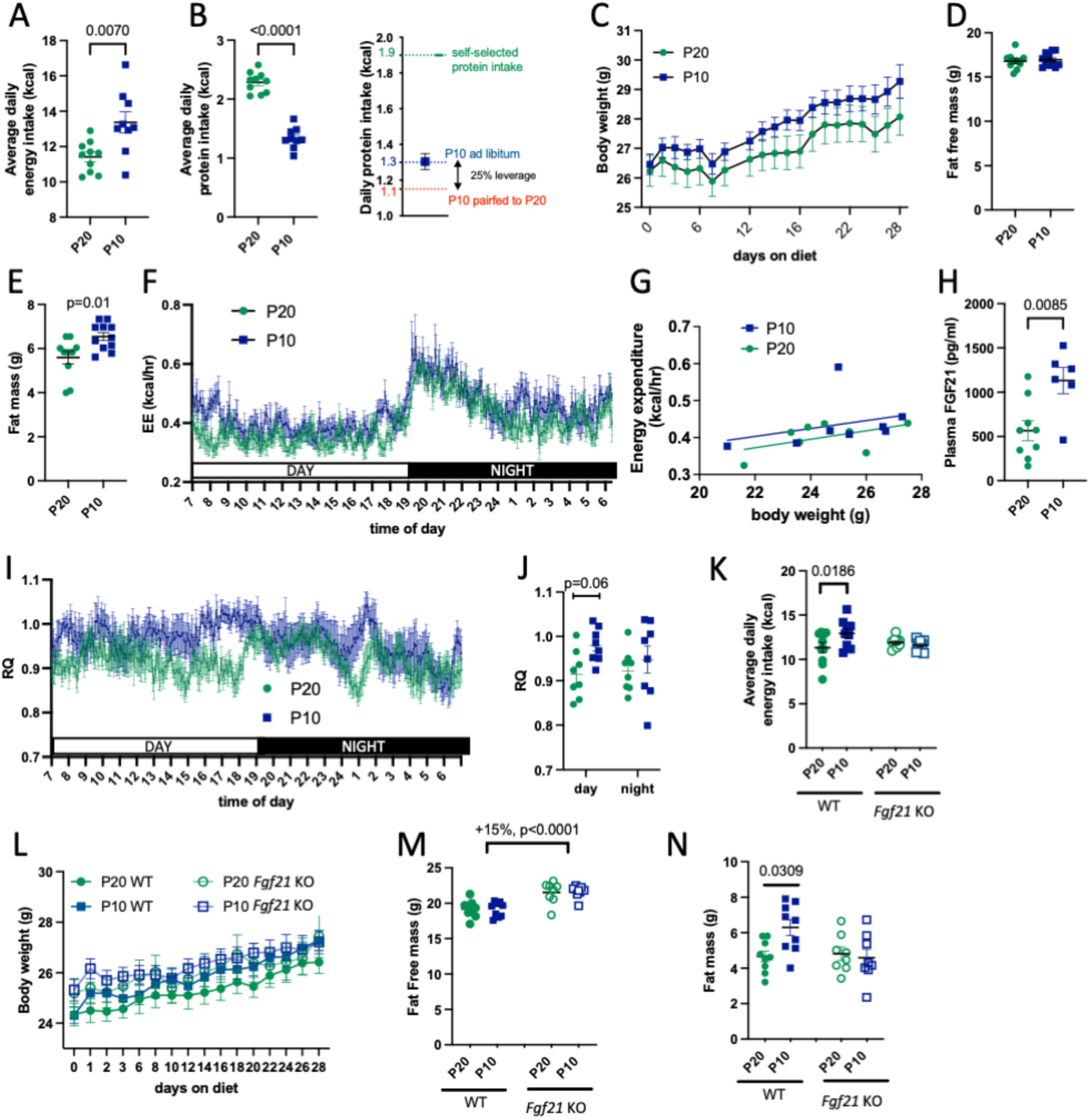
Mild protein restriction increases energy intake independently of changes in energy expenditure, increases adiposity but does not significantly affect weight gain in male mice. Energy intake (**A**), protein intake (**B**), body weight (C), fat free mass (**D**), fat mass **(E)**, energy expenditure (**F, G**), plasma FGF21 (**H**) and respiratory quotient (RQ, **I,J**) in wild type mice fed P10 or P20 at 22°C (n=9-10). Energy intake (**K**), body weight (**L**), fat mass (**M**) and fat free mass **(N)** in *Fgf21* KO mice and WT littermates fed P10 or P20 at 22°C (n=9-10). Data are means+/- sem.

The sustained hyperphagia in P10-fed mice in the absence of changes in metabolic rate suggests that mild protein dilution increases energy intake independently of energy expenditure under these conditions, and potentially independently of FGF21 signalling. However, FGF21 KO mice failed to develop hyperphagia or increased adiposity on the P10 diet (**Fig. 2K-2N**), indicating that FGF21 deletion interferes with the behavioural and metabolic adaptations to mild protein restriction. Of note, FGF21 KO mice defended a higher amount of lean mass (**Fig. 2M**), as previously reported^17^, suggesting altered baseline AA metabolism in this model.

### The hyperphagic response to mild protein restriction is blunted at thermoneutrality

At 22°C, the high energetic demand for thermoregulation might hinder the development of a body weight phenotype in mice fed the P10 diet despite their significant hyperphagia. To test this, we assessed maintained mice on the P10 and P20 diets at thermoneutrality (28°C). Unexpectedly, at 28°C the P10 diet only produced transient hyperphagia over the first 24h diet (**Fig. 3A**), after which energy intake remained comparable between P20 and P10-fed (**Fig. 3B**). As a result, daily protein intake decreased to an average of 1 kcal/day (**Fig. 3C**). Despite this drastic decrease in protein ingestion, body weight gain and lean mass did not change significantly between P10 and P20 mice under these conditions (**Fig. 3D, 3E**), and fat mass increased by 25% (**Fig. 3F**). Of note, plasma FGF21 levels increased significantly at 28°C with the P10 diet (**Fig. 3G**), but this increase failed to alter energy intake, suggesting that increased FGF21 signalling is not sufficient to produce hyperphagia during protein restriction.

**Figure 3:**
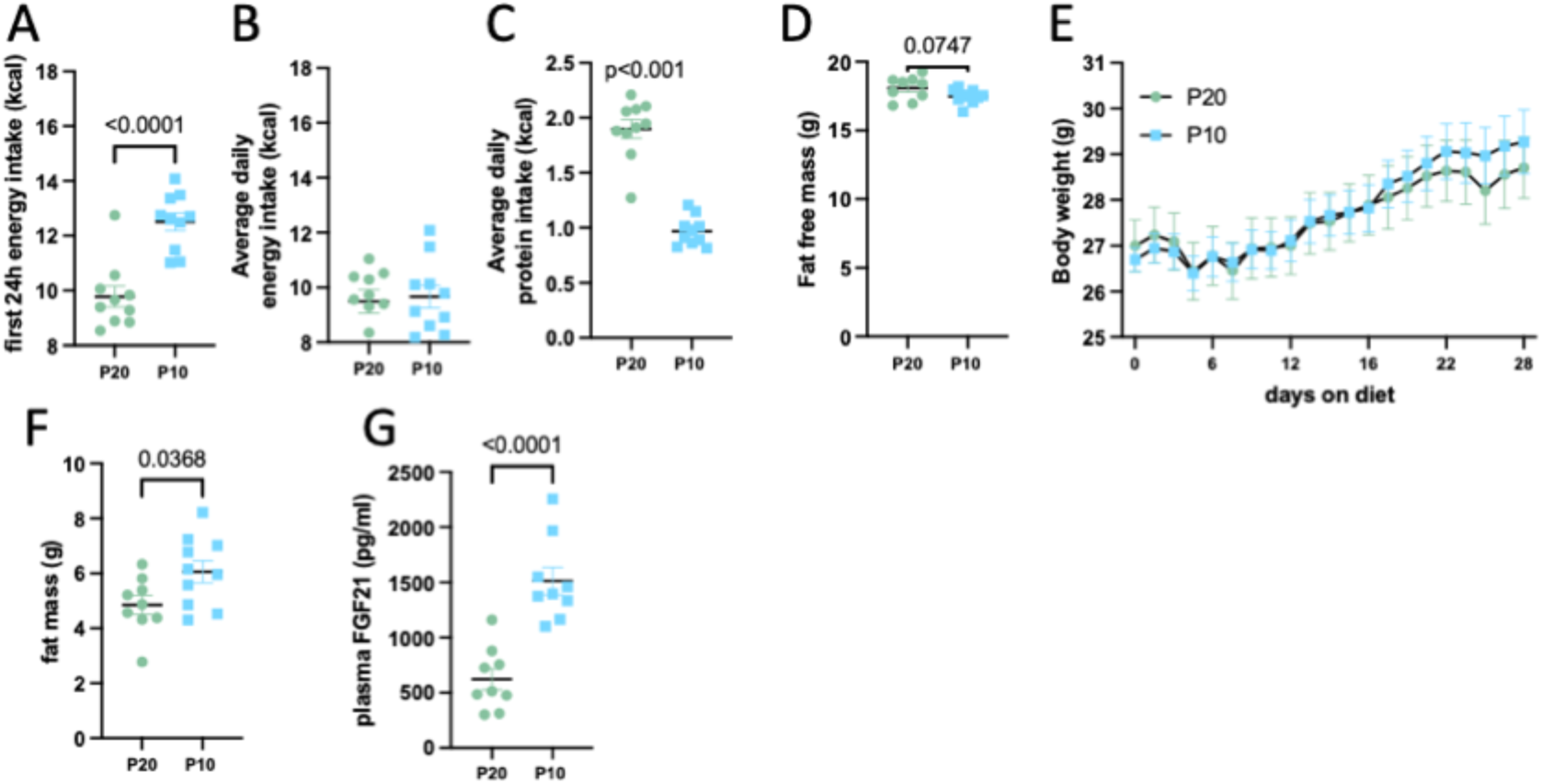
The hyperphagic response to mild protein restriction is blunted at thermoneutrality. Energy intake during the first 24h (**A**), average energy intake (**B**), average protein intake (**C**), body weight (**D**), fast free mass (**E**), fat mass (**F**) and plasma FGF21 (**G)** in mice fed P20 or P10 diet at 28 °C (n=9-10). Data are means+/- sem.

### Mild protein restriction produces protein preference at 22°C but not at 28°C

In addition to hyperphagia, protein restriction drives robust protein preference in mice maintained on a P5 diet, and this response requires central FGF21 signalling ^11,18,19^. Likewise, mild protein restriction with the P10 diet at 22**°**C produced protein preference and increased protein intake during a 24hr choice session where mice were offered ad libitum access to P5 and P45 pellets. P10 protein-restricted mice consumed less P5 (**Fig. 4A**) and more P45 (**Fig. 4B**) compared to mice maintained on P20, leading to a significant increase in P45 preference (**Fig. 4D**) and protein intake (**Fig. 4E**). Consistent with previous reports, *Fgf21* KO mice did not exhibit protein preference during protein restriction (**Fig. 4F-H**). Instead, P10-fed *Fgf*21 KO mice decreased rather than increased their intake of the P45 diet during the protein preference test (**Fig. 4I**), leading to decreased protein intake (**Fig. 4J**).

**Figure 4:**
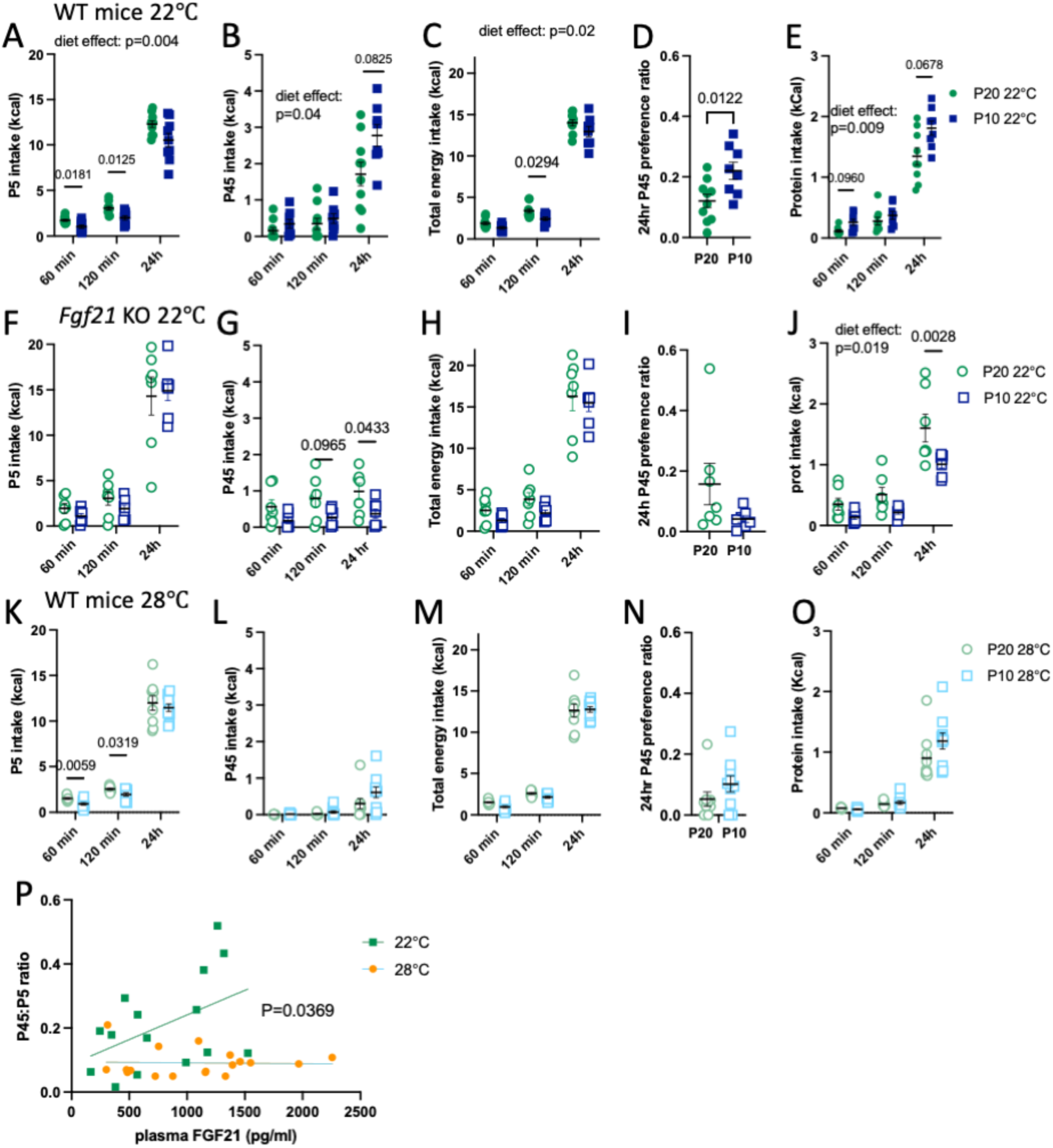
Mild protein restriction produces protein preference at 22°C but not at 28°C. Intake of P5 diet (**A, F, K**), and P45 diet (**B, G, L**), total energy intake (**C, H, M**), P45 preference ratio (**D, I, N)**, and total protein intake (**E, J, O**) during a 24h self-selection paradigm in WT (**A-E, K-O**) or *Fgf21* KO mice (**F-J**) maintained on P20 or P10 at 22**°**C (**A-E, F-J**) or 28**°** C (**K-O)**. (**P**) Correlation between plasma FGF21 levels and P45:P5 preference ratio in WT mice maintained on P20 or P10 at 22**°** C or 28**°**C. (**P**) Correlation between plasma FGF21 concentration and protein preference ratio in WT mice at 22**°** C and 28**°** C.

In contrast in mice adapted to thermoneutrality, P10 feeding did not produce preference for the protein-rich diet and did not increase protein intake during the choice test (**Fig. 4K-O**) despite the marked elevation in circulating FGF21 levels in P10-fed mice under these conditions (**Fig. 3G**). In fact, both P20-fed and P10-fed mice consumed marginal amounts of the P45 diet during the 24hr choice session at 28**°**C (**Fig. 4G**). Protein preference ratio significantly correlated with plasma FGF21 levels at 22**°**C but not at 28**°**C (**Fig. 4P**). Thus, increased FGF21 signalling is not sufficient to produce protein preference at thermoneutrality.

### Housing temperature determines the feeding and body weight responses to dietary protein dilution

The lack of hyperphagic response to mild protein restriction at thermoneutrality (**Fig. 3B**) suggests that thermoneutral housing blunts the adaptations to protein dilution. To further explore this, we used a diet containing 7% protein (P7), which still allows normal body weight and lean mass maintenance in male C57/Bl6J mice ^20^.

Mice fed the P7 diet at 22°C increased their daily energy intake by 20% (**Fig. 5A-B**), providing ∼ 1kcal protein per day (**Fig. 5C, S1A**), which was sufficient to defend their lean mass and body weight (**Fig. 5D, 5G**). Under these conditions, we observed a significant increase in adiposity (**Fig. 5H**), which was comparable to the response to P10-feeding at 22°C (**Fig. 2**). In sharp contrast, P7-feeding at 28°C only produced transient hyperphagia during the first 24h (**Fig. 5A**), after which energy intake was comparable between P20- and P7-fed mice (**Fig. 45**). Maintenance on P7 at 28°C produced significant weight loss (**Fig. 5E**) and loss of lean mass (**Fig. 5G**). Weight loss at 28°C occurred during the first 24h of P7 introduction, after which mice in fact defended a lower body weight at 1g below their baseline on average, which was accounted for exclusively by a loss in lean mass. These data confirm that thermoneutral conditions blunt the physiological adaptations to protein dilution and the mechanisms involved in the maintenance of body weight and lean mass set points. From these data we also conclude that a daily protein intake of ∼1 kcal per day (50% lower than the self-selected amount in macronutrient free choice conditions, **Fig. 1**) is sufficient to defend body weight and lean-mass, suggesting the recruitment of protein-sparing mechanisms.

**Figure 5:**
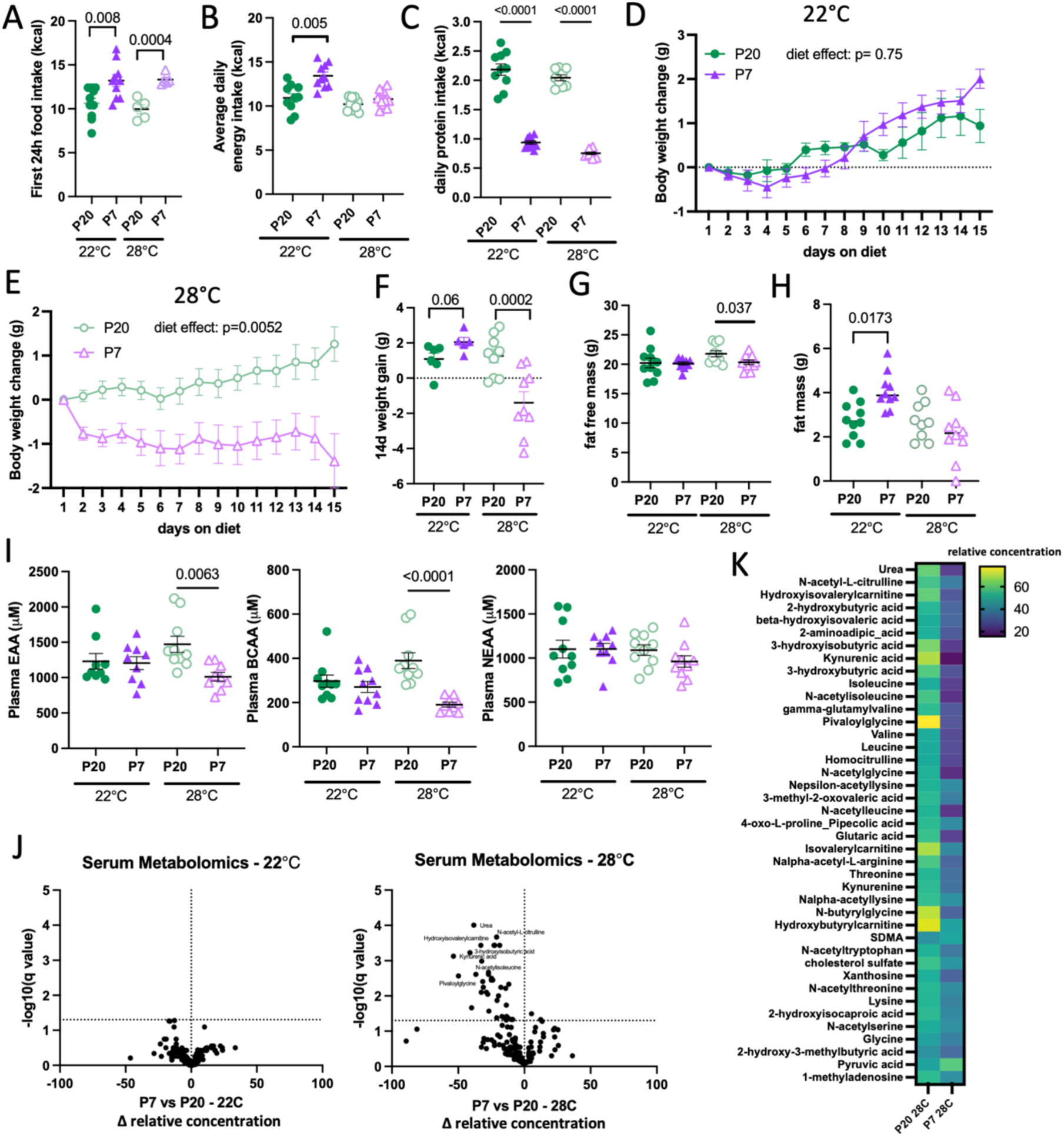
Housing temperature determines the metabolic and body weight consequences of dietary protein dilution. Food intake during the first 24h (**A**), average daily energy intake (**B**) average daily protein intake (**C**), body weight change (**D-F**), fat free mass (**G)** and fat mass **(H)** in mice fed P20 or P7 diets at 28°C or 22°C (n=9-10). (**I**) Interprandial plasma AA concentrations in mice maintained on the P20 and P7 diets at 22°C or 28°C (n=9-10). (**J**) Volcano plots of serum metabolome changes in P7-vs. P20-fed mice expressed as differences in relative concentrations against pooled calibrators, at 22°C and 28°C, and (**K**) heat map of significantly changed metabolites in serum of P7 vs. P20-fed mice at 28°C. Data are means +/- sem.

### Mice defend interprandial plasma EAA levels at 22°C but not at 28°C or in the absence of FGF21

The lack of change in lean mass and weight gain in mice fed the P7 diet at room temperature (**Fig. 5D, 5G**) despite a daily protein intake at 50% of the self-selected value suggests that mice can adapt well to mild protein restriction without having to reduce lean mass as a source of EAA. To determine the impact of mild protein restriction on circulating AA availability, we measured inter-prandial plasma AA levels in mice maintained on the P7 or P20 diets at room temperature or thermoneutrality. Remarkably, mice fed the P7 diet at 22°C maintained interprandial concentrations of circulating AA levels similar to those measured in control conditions (**Fig. 5I**). This was also the case for most AAs under more extreme dietary variations in protein intake, i.e. in mice fed either a 5% protein diet or a 45% protein diet for 7 days (**Fig. S1B-S1F**). The only significant changes were a 1.3-fold increase in BCAA levels and decreased alanine concentrations in P45-fed mice, the latter indicating increased gluconeogenesis.

**Figure S1:**
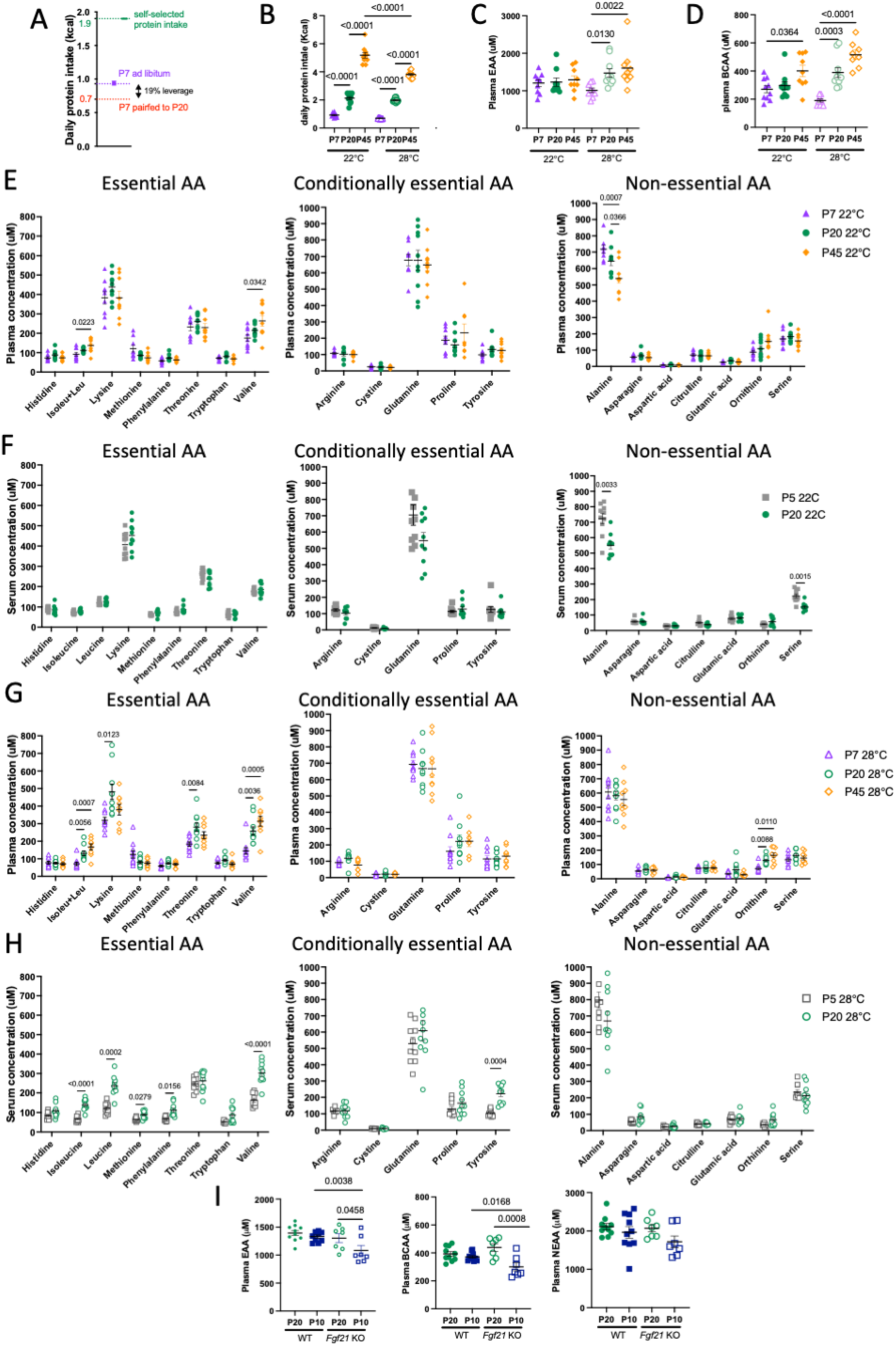
Protein intake in P7-fed mice at 22°C vs. target protein intake (**A**). Daily protein intake (**B**), interprandial plasma concentrations of EAA (**C**), BCAA (**D**), individual in mice maintained on P7, P20 or P45 diets for 1 week at 22°C (**E,** n=9-10) or 28°C (**F,** n=9-10). Interprandial serum concentrations of individual AAs in mice maintained on P5 or P20 diets for 1 week at 22°C (**G,** n=9-10) or 28°C (**H,** n=9-10). (**I**) Interprandial AA concentration in *Fgf21* KO mice and WT littermates following 7 days feeding with the P20 or P10 diets (n=7-10). Data are means +/- sem.

In contrast at 28°C, we observed a significant decrease in circulating levels of essential AAs and branched-chain AAs in P7-fed mice vs P20-fed controls (**Fig. 5I**). High protein feeding (P45) produced a further significant increase in circulating levels of branched-chain AAs (**Fig. S1G**), while severe protein restriction with the P5 diet reduced circulating concentrations of most EAAs (**Fig. S1H**). Thus, housing at thermoneutrality blunts the mechanisms buffering interprandial circulating AA levels.

Serum targeted metabolomics analysis confirmed these results and revealed a lack of change in circulating metabolites at 22°C in P7-fed mice compared to P20-fed mice (**Fig. 5J**). In contrast P7-fed mice at 28°C had a distinct serum metabolomic profile from control mice. Specifically, concentrations of most EAA and many AA-related metabolites were decreased (**Fig. 5J, 5K**).

Since lack of FGF21 signalling interferes with the behavioural and metabolic adaptations to protein restriction, we assessed the regulation of inter-prandial circulating AA levels in FGF21 KO mice maintained on the P20 or P10 diets. Unlike wild-type mice, FGF21 KO mice failed to maintain inter-prandial circulating levels of essential and branched-chain AAs during mild protein restriction at 22°C (**Fig. S1I**), further suggesting dysfunctional AA homeostasis in this model.

### Metabolome response to protein restriction

To gain insights into the mechanisms mediating the maintenance of circulating AA levels at 22°C, but not at 28°C, we performed targeted metabolomic analyses of liver, inguinal white adipose tissue (iWAT), epididymal white adipose tissue (eWAT), interscapular brown adipose tissue (BAT), gastrocnemius and soleus muscle, as well as two brain regions involved in nutrient sensing and peripheral metabolic control, i.e. the mediobasal hypothalamus (MBH) and dorso-vagal complex (DVC) in mice fed the P7, P20 and P45 diets at 22°C or 28°C for 7 days.

At 22°C, protein restriction with the P7 diet did not affect tissue EAA availability and produced only limited changes in tissue metabolites in skeletal muscle, MBH, DVC, iWAT and iBAT (**Fig. 6A**). Urea concentration was consistently lower in these tissues in P7-fed mice vs. P20-fed controls (**Fig. 6A**). Further, we found an increase in 1- and 3-methyl-L-histidine concentration in soleus muscle, MBH and DVC, a marker of protein breakdown ^21^ (**Fig. 6A**). Likewise, P45 feeding did not significantly change tissue EAA availability and upregulated urea concentration across most tissues, indicating a bidirectional control of tissue protein degradation in response to changes in dietary protein intake (**Fig. 6B**). Tissue levels of hydroxyvalerylcarnitine were consistently reduced in the soleus muscle, MBH, DVC, and iWAT in P7-fed mice, indicating a decrease in C5-acylcarnitine production from the BCAA L-leucine in these conditions (**Fig. 6A**). Strikingly, none of these responses occured in the liver in P7- or P45-fed mice (**Fig. S2B**), but nevertheless hepatic free AA levels were maintained across dietary conditions at 22°C (**Fig. 6B**).

**Figure 6:**
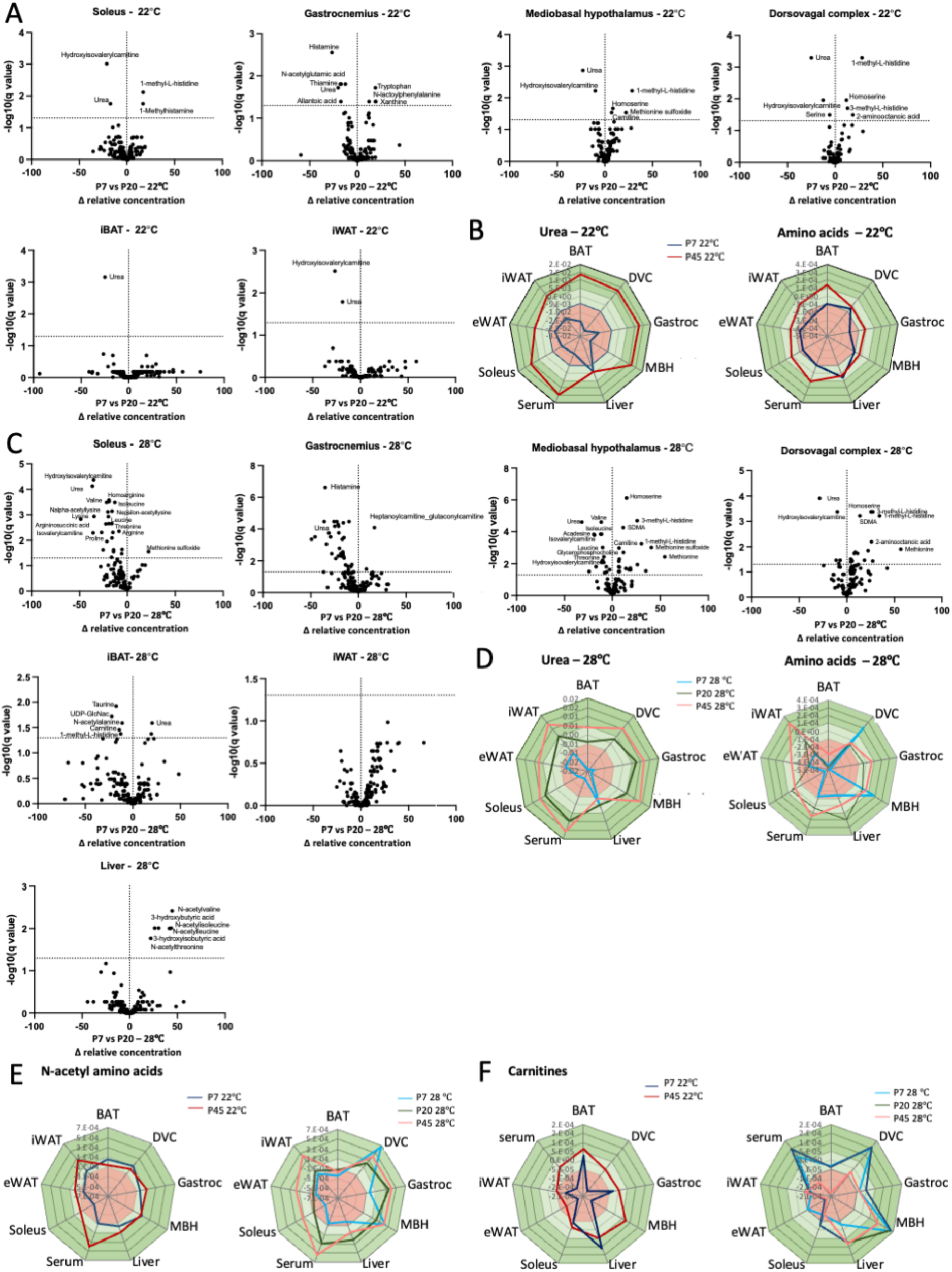
Metabolome responses to protein restriction. Volcano plots showing the difference in metabolite concentrations in mice fed the P7 vs P20 diets at 22**°**C (**A**) and 28**°**C (**C**), expressed as differences in relative concentrations against pooled calibrators. Error-normalised fold change in specific groups of metabolites against metabolite levels in the P20 group at 22**°**C ^24^ (**B,D-F**).

**Figure S2:**
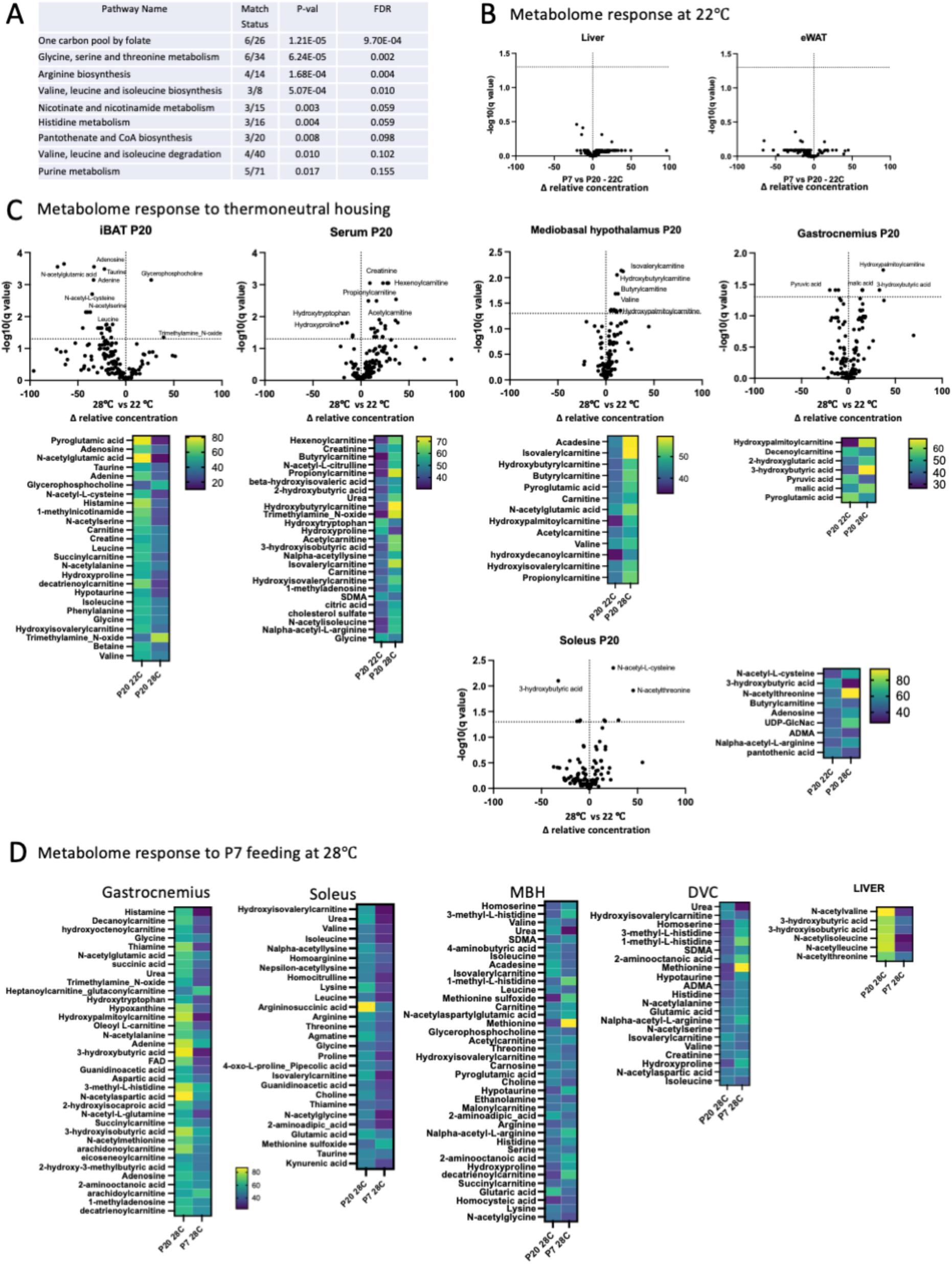
(**A**) Pathway analysis of top metabolites. Volcano plots showing the difference in metabolite concentrations in mice fed the P7 vs P20 diets at 22**°**C (**B**). Volcano plots showing the difference in metabolite concentrations in mice fed the P20 diet at 22**°**C vs 28**°**C and heat maps of significantly different metabolites (FDR<0.05) (**C**) Heat maps of significantly different metabolites in P7 vs. P20 mice at 28**°**C (FDR<0.05).

Housing at 28°C produced significant changes in metabolites levels in the iBAT, serum, skeletal muscle and MBH (**Fig. S2C**). In iBAT, we found decreased levels of EAA (e.g. BCAA), AA-related metabolites, purines (adenine, adenosine) and acylcarnitines (e.g. succinylcarnitine), consistent with decreased thermogenic activity ^22,23^. Parallel increases in BCAA-related metabolites and acylcarnitines were observed in the circulation, MBH and skeletal muscle (**Fig. S2C**), while the metabolomes of liver, eWAT and DVC were not altered. This suggests that decreased iBAT thermogenic activity at 28°C leads to a spillover of acylcarnitines and AA-related metabolites to the circulation, skeletal muscle and MBH.

P7 and P45 feeding at 28°C produced similar changes in urea levels as seen at 22°C (**Fig. 6D**) but despite this, EAA levels were reduced in serum, soleus, MBH and DVC during P7 feeding (**Fig. S2D**, 6**J-K**), and increased in serum and iWAT during P45 feeding (**Fig. 6D**). In P7-fed mice, N-acetyl AAs, AA-related metabolites, purines and acylcarnitines were significantly reduced in skeletal muscle, MBH and DVC (**Fig 6C, S2D**). We found only a few changes in the liver of P7-fed mice at 28°C, specifically decreased levels of BCAA related and N-acetyl AAs (**Fig. 6D**). Across conditions, serum N-acetyl AAs levels reflected changes in dietary protein intake both at 22°C and 28°C, with stable tissue levels at 22°C but not at 28°C (**Fig. 6E),** hinting to a potential role for N-acetyl AAs in buffering changes in dietary protein intake and dysregulated at 28°C. Likewise, serum, DVC, MBH and eWAT carnitine levels reflected dietary protein intake, but this regulation was blunted in the serum, MBH and DVC at 28°C (**Fig. 6F**)

### iBAT BCAA catabolism buffers circulating BCAA levels at 22°C but not at 28°C

BCAA-related metabolites, N-acetyl AAs and acylcarnitines were among the top metabolites altered in during dietary protein restriction. Therefore, we next measured the expression of genes involved in BCAA catabolism (*Bcat2, Bckdha, Bckdhb, Bckdk, Hibch, Slc25a44*), carnitine metabolism (*Cpt1a*, *Cpt1b*, *Cpt2*, *Crat*, *Crot*) , and aminoacylation (N-acetyltransferase 8, *Nat8* and Aminoacylase 1, *Acy1*) in skeletal muscle depots (gastrocnemius and tibialis anteralis), liver and iBAT.

We found only modest changes in expression of these genes in skeletal muscle depots in response to protein restriction at both housing temperatures (**Fig. 7A, 7B**). We found decreased expression of genes involved in carnitine metabolism at 28°C, more pronounced in P7-fed mice, consistent with reduced acylcarnitine levels during protein restriction at thermoneutrality (**Fig 7F, S2D**). *Nat8* expression was increased at 28°C in both P20- and P7-fed mice in the gastrocnemius muscle.

**Figure 7:**
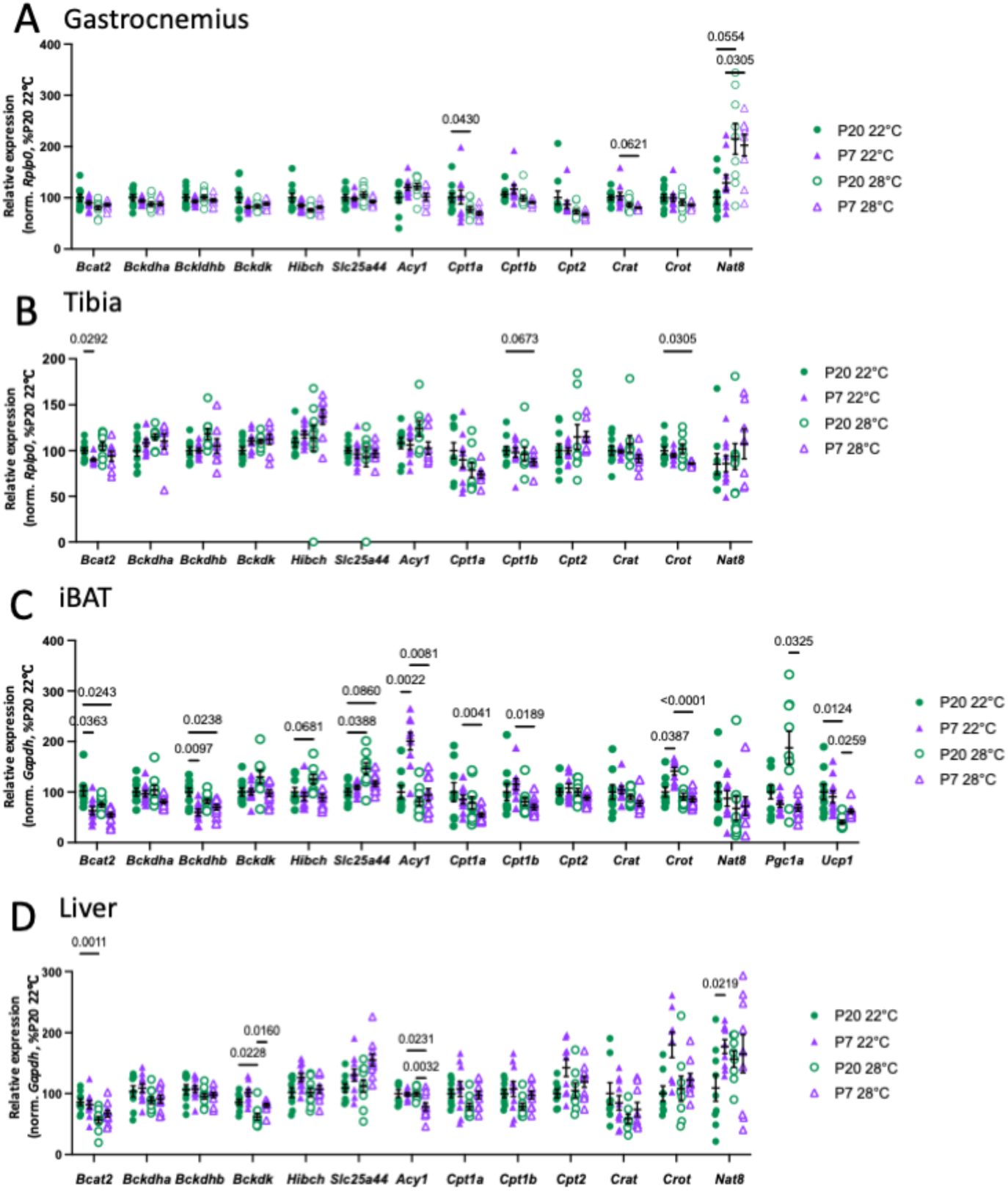
Gene expression analysis in the gastrocnemius muscle (A), tibialis anterior (B), iBAT (C) and liver (D) in mice fed the P7 vs P20 diets at 22**°**C and 28**°**C . Data are means ±sem.

In iBAT, protein restriction at 22°C decreased the expression of genes involved in BCAA catabolism (*Bcat2* and *Bckdhb*) and increased the expression of *Acy1* (**Fig. 7C**). These changes were blunted at thermoneutrality. iBAT *Crot* expression increased with P7-feeding at 22°C, indicating a switch in substrate utilization for thermogenesis. Adaptive changes to thermoneutral housing in iBAT of control mice included increased expression of the mitochondrial BCAA transporter *Slc25a44*^22^ *a*nd decreased expression of *Ucp1*, but these changes were blunted in P7-fed mice.

In the liver, protein restriction at 22°C did not change the expression of BCAA catabolic genes but increased Nat8 expression (**Fig. 7D**), suggesting a recruitment of AA N-acetylation pathways to maintain hepatic AA levels during protein restriction. At 28°C, protein restriction decreased *Acy1* expression, suggesting and additional mechanism to buffer hepatic AA levels using the acylated AA pool. Thermoneutral housing decreased *Bckdk* expression, suggesting increased hepatic BCAA catabolism in these conditions.

## Discussion

This study provides evidence that protein intake is controlled independently of energy intake in male mice allowed to self-select the protein-carbohydrate ratio of their diet, and maintained at around 1.9 kcal per day. This supports the contribution of protein-sensing pathways in the control of food intake and raises the question of how these pathways are integrated with energy-sensing pathways and the regulation of energy homeostasis, particularly when dietary protein availability is low. The protein leverage hypothesis predicts that under such condition, the need to obtain a fixed amount of protein (our data suggest this would be 1.9kcal in adult male mice) would be prioritised over the regulation of energy homeostasis, driving hyperphagia and weight gain. We tested this prediction using low protein diets containing 10% or 7% protein. These diets are compatible with normal body weight maintenance, unlike more severe protein restriction paradigms ^11,12^. With both diets, we modelled some key aspects of the protein leverage hypothesis including hyperphagia and protein-specific appetite, but daily protein intake remained significantly lower than 1.9 kcal protein. We measured an increase in adiposity with both diets, but no significant change in body weight. This was also the case in mice housed at thermoneutrality, indicating that the regulation of energy homeostasis is prioritised over the maintenance of dietary protein intake under these conditions.

Our model provides insights into the mechanisms engaged during the adaptation to protein restriction without loss of lean mass or interruption of normal growth. Under these conditions, protein restriction produced hyperphagia in the absence of changes in energy expenditure, indicating that hyperphagia can be a primary response to protein deficit. This contrasts with previous finding suggesting that hyperphagia occurs secondary to hypermetabolism in protein restricted mice ^25^. The rationale for testing the feeding and metabolic consequences of mild protein restriction at thermoneutrality was to determine whether alleviating the chronic thermal stress at 22°C might potentiate the consequences of mild protein restriction on adiposity and body weight gain. Against our prediction, P10-fed and P7-fed mice were unable to mount the appropriate adaptive behavioural responses to protein restriction at 28°C, resulting in loss of lean mass and decreased circulating levels of EAA. This occurred despite elevated FGF21 levels, supporting the conclusion that increased FGF21 signalling is not sufficient to produce hyperphagia and protein preference. In addition, maintenance on the P10 diet produced a significant increase in adiposity, which is maintained at thermoneutrality in the absence of hyperphagia. Eventually, increased adiposity in during mild protein restriction might lead to obesity and associated metabolic dysfunctions over time, yet this occurred independently of energy intake at thermoneutrality, thus through mechanisms distinct from protein leverage.

One major finding of this study is that circulating and tissular concentrations of free EAA are tightly regulated during protein restriction and to a large extend during high-protein feeding when mice are maintained at 22°C, including in the skeletal muscle, first site of AA oxidation ^26^, and the liver, proposed to mediate dietary protein sensing and the associated behavioural and metabolic responses ^27^. In protein restricted mice, this regulation occurs despite protein intake being below the self-selected value of 1.9kcal, indicating the recruitment of AA buffering mechanisms. We conclude that homeostatic mechanisms efficiently maintain circulating AA availability despite wide variations in dietary protein intake.

Our metabolomic analyses provide insight into the mechanisms allowing the maintenance of AA homeostasis. Reduced urea concentrations during protein restriction across most examined tissues indicates that decreased AA oxidation is a ubiquitous mechanism sparing AAs in tissues. Strikingly, although the urea cycle occurs mainly in the liver ^28^, hepatic urea concentration did not change during either low or high protein feeding, highlighting the role of extra-hepatic nitrogen disposal mechanisms in the regulation of AA homeostasis. We also observed a wide-spread decrease in acylcarnitine synthesis from BCAA metabolites, as suggested by changes in hydroxyisovalerylcarnitine levels. We identified iBAT AA catabolism and carnitine synthesis as the main pathways regulated by protein restriction at 22°C, through the downregulation of *Bcat2*, *Bckdh* and upregulation of *Crot* expression. In addition, elevated iBAT *Acy1* expression identifies a role for aminoacylation and release of free AA from the N-acyl-L-AA pool to maintain free AA levels.

The role of the iBAT is further highlighted by the disruption of AA homeostasis at 28° C, evident even in P20-fed mice (elevated AA and acylcarnitine levels in the circulation and the brain), and exacerbated during high or low protein feeding. iBAT is the main tissue recruited for adaptive thermogenesis, together with skeletal muscle, recruited for shivering thermogenesis, and the brain, which regulates body temperature ^29^. These tissues exhibit the most metabolome changes during protein restriction at 22°C, and following adaptation to thermoneutral housing on the control diet. However, metabolomic data indicate that the downregulation of AA oxidation and acylcarnitine synthesis at thermoneutrality occurs specifically in iBAT. In addition, P7-induced gene expression changes for reduced BCAA catabolism and aminoacylation are blunted at 28°C. This is consistent with the previously reported role of iBAT as a site for BCAA clearance through thermogenesis in mice and humans ^22^ . We conclude that adaptations in iBAT AA utilisation are required for the maintenance of AA homeostasis in response to changes in dietary protein intake, and propose that the loss of this buffering mechanism at 28°C accounts for the failed maintenance of AA homeostasis at 28°C.

The liver differs from other tissues in its adaptations to changes in protein intake and ability to maintain free AA concentrations even at 28°C, when systemic AA homeostasis is compromised. The lack of change in hepatic metabolome during protein restriction is consistent with a previous report, where severe protein restriction did not perturb hepatic EAA concentrations, independently of ATF4 signalling ^30^. This contrasts however, with liver transcriptomic data indicating downregulation of hepatic arginine biosynthesis, involved in urea production, and glycine/serine/threonine metabolism in protein restricted mice ^20^. Also, our data do not indicate a reduction in hepatic BCAA oxidation at 22° C, unlike previous reports ^31^. Here, we identify distinct mechanisms recruited for the maintenance of hepatic AA levels. At 22° C, upregulation of hepatic *Nat8* expression highlights a role for N-acetyl AAs to buffer hepatic AA levels against variations in dietary intake. At 28° C, this mechanism is blunted and instead, AA catabolism is reduced, as suggested by the increased expression of *Bckdk* in P7-fed mice at thermoneutrality. In addition, P7 feeding at 28C reduced expression of hepatic *Acy1*, which might contribute to the failed maintenance of circulating AA under these conditions.

Our results reveal the high sensitivity of the MBH and DVC metabolism to protein restriction at 22°C. These two brain sites are important AA sensing regions ^32,33^, and our data suggest that local sensing of AA metabolites might also be relevant. Further work is needed to identify brain AA sensing mechanisms and their role in systemic AA homeostasis. This contrasts with the current view according to which the brain is indirectly involved in the adaptations to protein restriction, in response to hepatic FGF21 signaling ^11^. Our data suggest that FGF21 is not the sole signal mediating the behavioural adaptations to protein restriction. FGF21 has been shown to be induced by protein restriction^34^ and is elevated even during mild protein restriction here, but not sufficient to produce hyperphagia at thermoneutrality. Others have reported discrepancies in the FGF21-centric model of protein restriction, for example showing that females exhibit the expected adaptation to protein restriction despite sowing no change in circulating FGF21 levels ^20^. We propose that the lack of response to protein restriction in FGF21 KO^11^ or even UCP1 KO^25^ mice might be in fact confounded by their suboptimal ability to adaptively recruit thermoregulatory pathways, which has been shown during cold stress for example ^35^ or in response to adrenergic stimulation ^36^. More work is needed to fully understand the mechanism generating the hyperphagic response to protein restriction. It might also rely on the downregulation of iBAT activity at 28°C. Previous work supports a role of the iBAT in relaying information on nutritional status to the brain, for example through secretin.

Downregulation of the iBAT-brain appetite-regulating axis at thermoneutrality. Consistent with this hypothesis, the secretion of secretin is blunted at thermoneutrality and in UCP1 KO mice ^37^.

A question that arises from our results is the evolutionary fitness of such mechanism that allows optimal physiology (maintenance of AA homeostasis) in suboptimal environmental conditions (housing below the thermoneutral point). Considering this question, it’s important to note that mice evolved a unique thermoregulatory strategy driven by their high metabolic capacity ^38^. Although mice prefer warmer temperature, one can argue that they did not evolve in environments at 28C, and thus some physiological function might be best maintained below thermoneutrality. Humans do not have a thermoneutral point, and burn calories for thermoregulations across a wide range of ambient temperature ^39^. Thus, the relevance of our findings to human physiology should be evaluated in future studies.

In summary, our data provide new insights into the mechanisms recruited to produce the behavioural and metabolic responses to protein restriction, and the maintenance of AA homeostasis. Mild protein restriction might help characterise the central and peripheral mechanisms engaged in response to conditions of reduced dietary protein intake in which organisms successfully adapt to maintain normal physiology and AA homeostasis. Obesity and insulin resistance associate with dysregulations of AA metabolism. Future work should examine the contribution of BCAA buffering mechanisms identified here in the pathogenesis of metabolic diseases

## Methods

### Lead contact

Further information and requests for resources and reagents should be directed to and will be fulfilled by the Lead Contact, Clemence Blouet

### Materials availability

This study did not generate any new reagents

### Data and code availabilty

All data are available in the main text or supplementary materials.

### Animals

All animal experiments were performed in accordance with the UK Home Office regulations under the Animals (Scientific Procedures) Act (1986) and with the approval of the University of Cambridge Animal Welfare and Ethics Review Board (AWERB). Animals were group-housed in a specific pathogen free facility and maintained on a standard 12-hour light/dark cycle (lights on 7:00-19:00) with ad libitum access to water and food unless otherwise stated. For experiments performed at different housing temperatures, mice were maintained in temperature-controlled cabinets. All experiments were performed on male C57BL/6J mice obtained from Charles Rivers Laboratories and aged 9-wks old at the beginning of the experiments. *Fgf21* KO mice (B6N; 129S5-Fgf21tm1Lex/Mmucd) were generated on a C57BL/6N background using sperm obtained from MMRRC.

### Dietary manipulations

All mice were adapted to single housing and maintained on the P20 control diet for a week before being randomly allocated to an experimental diet. Before exposure to new diets, mice were allowed to taste new diets for 15 to 30 min or 3 consecutive days to avoid neophobia. All experimental diets were manufactured by Research Diets (Supple. Table 1).

### Ghrelin challenge

Mice were scruffed daily for a week to become acclimatised to ip injection paradigm. On the experimental day, mice received an ip injection 10ug/kg 1h before dark onset, and food intake was measured over 24hr.

### Macronutrient choice

Mice were presented with both the P5 and P45 diet 1h before dark onset and food intake was measured over 24hr.

### Body composition analysis

Body composition was analysed in conscious, free moving animals by time domain nuclear magnetic resonance (TD-NMR) using a MiniSpec LF90 Whole Body Composition Analyser (Bruker).

### Indirect calorimetry

Mice were single-housed for at least one week prior to housing in Promethion High-Definition Multiplexed Respirometry Cages (Sable Systems International) for 48 hours, during which time energy expenditure and activity data was collected. During analysis the first 24 hours of each run was discarded to allow for acclimatisation of mice to the calorimetric cages.

### Plasma analysis

Whole blood was collected from the tail-vein of live mice with a heparin-coated capillary (for FGF21 measurement), or terminally into an EDTA-treated tube (Sarstedt) via left ventricle cardiac puncture from mice under anaesthesia following an intraperitoneal injection of 100 mg/kg sodium pentobarbitone (for AA profiling).

Plasma FGF21 was analysed using a Quantikine ELISA kit (R&D Systems, product code MF2100) following the manufacturer’s instructions. This was performed by the Cambridge MRC MDU Mouse Biochemistry Laboratory and human assays were provided by the NIHR Cambridge BRC Core Biochemical Assay laboratory (CBAL).

For plasma AA analysis, 5 µl of plasma sample was added with mixed isotope-labelled AA internal standards, then extracted by protein precipitation using formic acid and isopropyl alcohol followed by derivatisation using AccQ-Tag Ultra Derivatising Kit (Water 186003836). Derivatized AA standards in 0.1% formic acid was prepared and analysed along with the extracted samples. Liquid chromatography and tandem mass spectrometry was performed on the Shimadzu Nexera X2 Liquid chromatography system linked to Sciex 6600 Triple Quad mass spectrometer. 19 proteinogenic AAs (except glycine due to protocol limitations) plus citrulline, orthenine and taurine were analysed. This was performed by the Pharmacokinetics and bioanalysis core facility of the Cancer Research UK Cambridge Institute.

### Targeted metabolomics

Targeted metabolomics was performed on serum, liver, interscapular brown adipose tissue (iBAT), epididymal white adipose tissue (eWAT), inguinal adipose tissue (iWAT), soleus and gastrocnemius muscle samples, and microdissected mediobasal hypothalamus and dorsovagal complex samples using a liquid chromatography with tandem mass spectrometry (LC-MS/MS) approach, as described in ^40,41^. Briefly, plasma samples were frozen immediately after collection and then thawed in an ice bath prior to LC-MS/MS analysis. Two extraction protocols (methods A and B) were used to collect metabolomics data. For method (A), 150 μL of pre-chilled acetonitrile-methanol (1:1 v/v) solvent mixture was added to 25-μL plasma aliquot. For method (B), 150 μL of pre-chilled methanol-water (4:1 v/v) solvent mixture was added to a second 25-μL plasma aliquot. Samples were mixed thoroughly and incubated at −20°C overnight. Both extraction solutions were spiked with a mix of stable labelled internal standards. After overnight extraction, the samples were centrifuged for 15 minutes at 14000 g at +4°C. A pooled calibrator sample was created for each method by combining an aliquot of supernatant from all the extracted samples together. The pooled calibrator was serially diluted in the same extraction solvent and used to create a calibration curve (100% [pooled calibrator], 75%, 50%, 30%, 15%, 10%, and 5%) for relative quantitation and batch-to-batch data normalization. Individual samples were further diluted at 1:1 (v/v) ratio with each extraction solution, transferred into a new 96-well plate, and injected for LC-MS/MS analysis.

Tissue samples were flash frozen in liquid nitrogen immediately after collection and stored at −80 °C. Frozen tissue samples were pulverized using a tissue pulverizer and kept frozen until LC-MS/MS analysis. Two aliquots of pulverized tissue were weighted for each sample to run methods A and B independently. Tissue samples were extracted and processed as described above. Extraction solutions were added to reach a final concentration of 100 mg/mL.

Data were acquired using a Shimadzu Nexera X2 UPLC system coupled to an AB Sciex 6500+ triple quadrupole mass spectrometer equipped with an electrospray source. For method A, a Waters Acquity BEH Amide 100 mm x 2.1 mm, 1.7 mm particle size, column was used; for method B, a Waters XSelect HSS T3 C18 100 mm x 2.1 mm, 1.8 mm particle size, column was used. Both columns were maintained at +40C. Elution solvents for both methods were 10 mM ammonium formate adjusted with 0.1% formic acid (solvent A) and 0.1% formic acid in acetonitrile (solvent B). Data were acquired using scheduled multiple reaction monitoring mode with polarity switching. A total of 250 polar metabolites were targeted. For most of the metabolites, identification was supported by 1 qualifier ion monitored in addition to a quantifier ion. Two qualifier ions were monitored to resolve interferences amongst few metabolites (data not shown). Peak areas were integrated using the AB SCIEX MultiQuant 3.0.2 software. Only analytes detected at the lowest pooled quality control calibrator with signal-to-noise >3 and detected in more than 75% of the individual samples were quantified. Relative quantitation to the pooled calibrators was achieved using a linear regression model. Metabolite areas were normalized to internal standard area responses. For analytes with no matching stable labelled internal standards, the optimal choice (picked amongst the internal standards monitored under the same polarity and within the same assay) was the internal standard giving the minimum root mean square error for the pooled calibrators and best linear fit. Individual sample values falling below the limit of detection were imputed to 1/2 times the lower calibrator.

For statistical analysis, the metabolomics data were regressed against a pool reference sample. The reference sample was made by pooling together equal amounts of extracts for each sample in the batch, as previously described ^41^. A calibration curve was built from that pool by serial dilution (down to 5%, so a 20 fold dilution) into solvent containing internal standards. The signal from each sample was calculated as a percentage of the pool signal. Differences against 20% protein at 22C were used to generate the spider plots comparing all conditions. These were calculated using error-normalised fold change to describe differences, which is described in detail in ^42^. In addition, differences between the P20 and P7 diet at 22C or 28C were analysed using one-way ANOVA with Dunnett’s post hoc test was performed to assess significance (p < 0.05) of between-group data distributions for individual metabolites.

### RNA isolation, cDNA synthesis and qPCR

Total RNA of snap-frozen tissues was extracted in TRIzol reagent (Thermo Fisher Scientific) using Lysing Matrix D homogenisation tube and Fastprep 24 Homogenizer (MP Biomedicals). RNA was extracted using isopropanol phase separation and RNA precipitation with 70% ethanol. Precipitated RNA was then subject to a clean-up step with Qiagen RNeasy Mini Kit as per the manufacturer’s instructions. cDNA synthesis of 1 μg RNA was performed using the High-Capacity cDNA Reverse Transcription Kit with random hexamer primers (Thermo Fisher Scientific). qPCR was performed using the SYBR Green PCR Master Mix for SYBR Green reactions (Thermo Fisher Scientific) on a QuantStudio 5 Real-Time PCR System (Thermo Fisher Scientific). The average of technical duplicates of a biological sample was used for quantification.

Relative gene expression was quantified using the -ΔΔ threshold cycle (Ct) method. *Rplp0* was used as the reference gene for analysis of tibialis anterior and gastrocnemius muscles; *Gapdh* was used for liver and iBAT measurement. The specificity of all SYBR Green primers were validated with melting curve analyses. All qPCR primers were designed to target the exon-exon boundary or separated by a long intron to exclude amplifying genomic DNA; and were synthesised by Merck. Following qpcr primers were used:

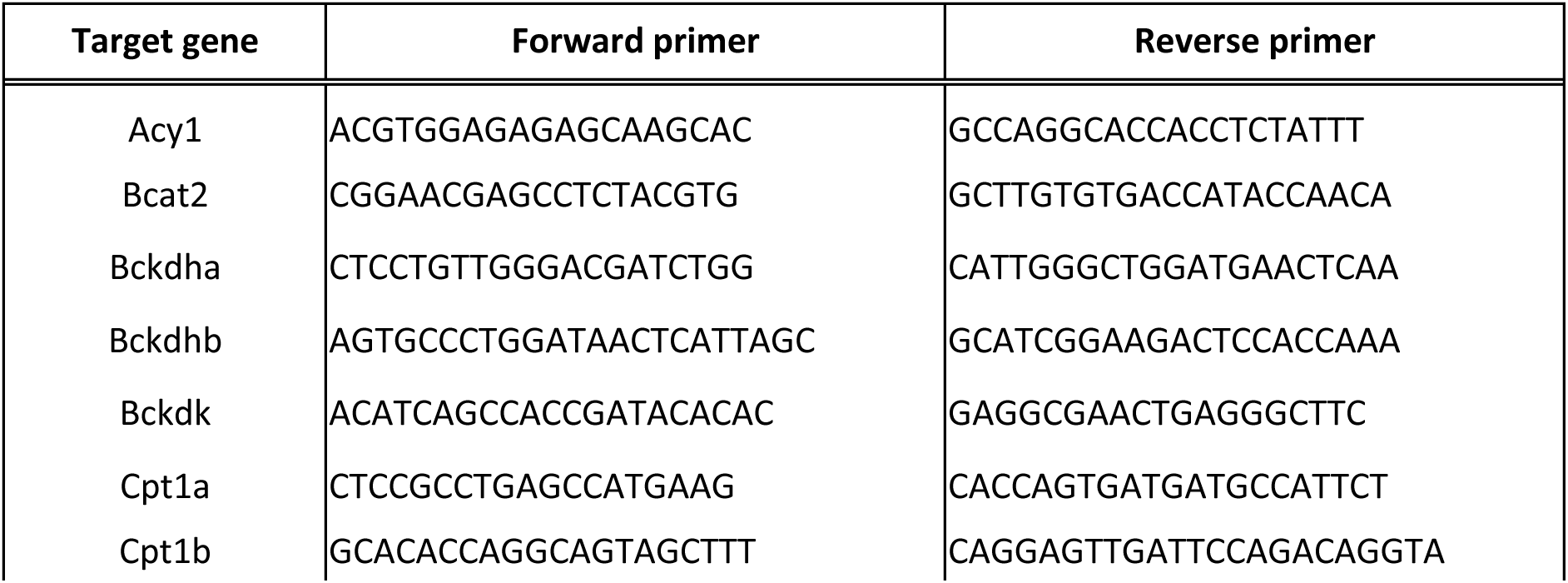
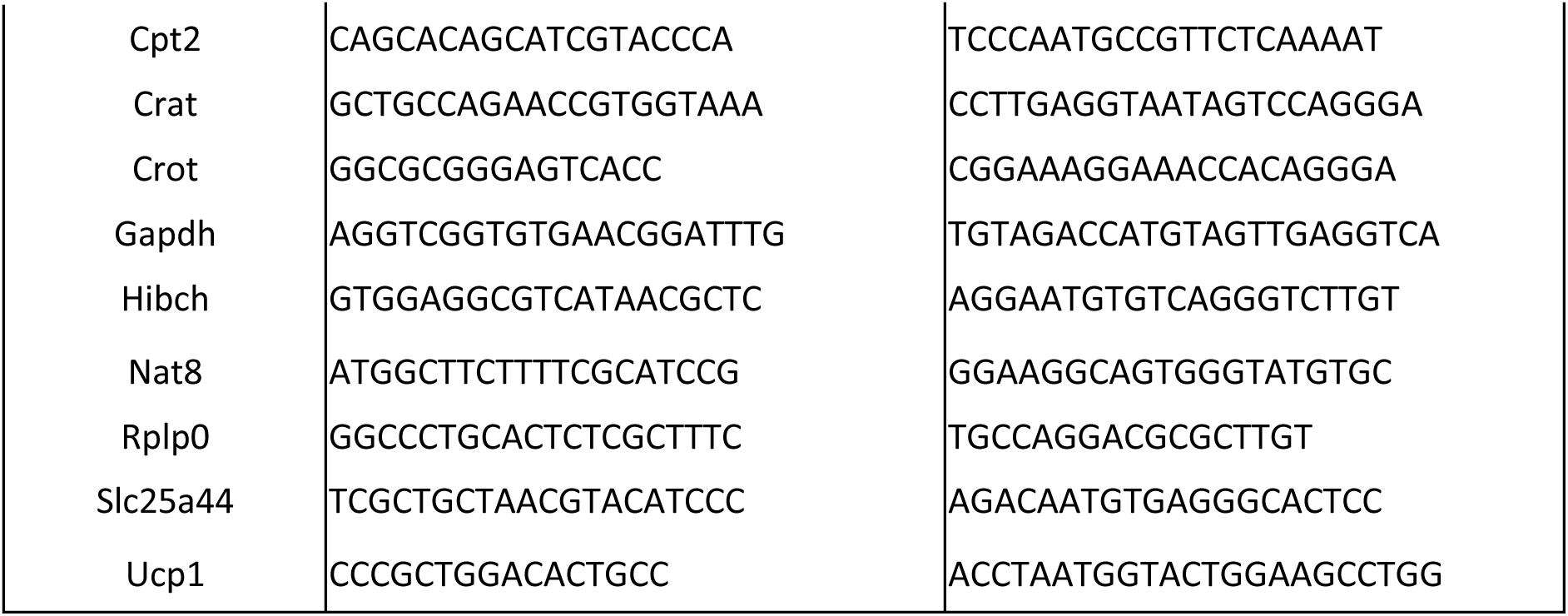

### Statistical analysis

All data visualisation and statistical analysis was performed in Prism 9 Software (GraphPad), except for metabolomics analyses (see details above). For all statistical tests, an α risk of 5% was used. All kinetics were analysed using repeated-measures two-way ANOVAs and adjusted with post hoc tests. Multiple comparisons were tested with one-way ANOVAs and adjusted with Tukey’s post hoc tests. Single comparisons were made using two-tail Student’s t tests.

## Data availability

### Lead contact

Further information and requests for reagents may be directed to and will be fulfilled by the Lead contact, Clemence Blouet

### Materials availability

This study did not generate new unique reagents.

### Data and code availability

**-** Data used to generate the figures in this paper will be shared by the lead contact upon request.
**-** This paper does not report original code.
**-** Any additional information required to reanalyze the data reported in this paper is available from the lead contact upon request.

## Acknowledgments

We thank the Histopathology, Imaging, Lipidomic, Disease Model Cores, in addition to the Core Biochemical assay Laboratory at the Institute of Metabolic Sciences for their contributions. We also thankthe CRUK Cambridge Institute Pharmacokinetics and Bioanalysis Core facility.This work was supported by a Medical Research Council grant (MR/S011552/1; CB), Medical Research Council Metabolic Disease Unit Grants (MC_UU_00014/5, MRC_UU_00014/1) and (MRC_MC_UU_12012/5) and a Wellcome Trust Strategic Award (208363/Z/17/Z).

## Author contributions

AT, SB, SV and CB designed the experiments. AT, SB, SL, AP, LP and CB executed experiments. AT, SB, CB and AK analyzed the data. AT, SV, TC and CB wrote the manuscript.

