## Supplementary material for "Adaptive thermogenesis reprograms the behavioural and metabolic responses to protein restriction for successful amino acid homeostasis": Supplemetal table 1

Supplementary Table 1: Diet composition

| | P5 | P7 | P10 | P20 | P45 |
|---|---|---|---|---|---|
| **Ingredient** | gm | gm | gm | gm | gm |
| **Casein** | 51.2 | 75 | 109 | 223 | 510 |
| **L-Cystine** | 3.9 | 3.7 | 3.5 | 3.5 | 3.5 |
| **DL-Methionine** | 3.61 | 3.1 | 2.25 | 0 | 0 |
| **Corn Starch** | 484 | 466.5 | 509.2 | 326.6 | 156.5 |
| **Maltodextrin 10** | 150 | 150 | 75 | 150 | 75 |
| **Sucrose** | 107.07 | 107.07 | 107.07 | 107.07 | 107.07 |
| **Cellulose** | 50 | 50 | 50 | 50 | 50 |
| **Soybean Oil** | 88.9 | 88.9 | 88.9 | 88.9 | 88.9 |
| **tBHQ** | 0.014 | 0.014 | 0.014 | 0.014 | 0.014 |
| **Mineral Mix S10022G** | 0 | 0 | 0 | 0 | 0 |
| **Mineral Mix S10022C** | 3.5 | 3.5 | 3.5 | 3.5 | 3.5 |
| **Calcium Carbonate** | 8.26 | 8.87 | 9.72 | 10 | 12.35 |
| **Calcium Phosphate, Dibasic** | 6 | 5.15 | 3.96 | 3.4 | 0 |
| **Potassium Citrate, 1 H20** | 0 | 0 | 0 | 3 | 8 |
| **Potassium Phosphate, Monobasic** | 10.11 | 10.11 | 10.11 | 6.31 | 0 |
| **Sodium Chloride** | 2.59 | 2.59 | 2.59 | 2.59 | 2.59 |
| **Vitamin Mix V10037** | 10 | 10 | 10 | 10 | 10 |
| **Choline Bitrartrate** | 2.5 | 2.5 | 2.5 | 2.5 | 2.5 |
| **FD&C Yellow Dye #5** | 0.025 | 0.05 | 0 | 0 | 0 |
| **FD&C Red Dye #40** | 0 | 0 | 0.025 | 0.05 | 0 |
| **FD&C Blue Dye #1** | 0.025 | 0 | 0.025 | 0 | 0.05 |
| **Total** | 978.58 | 987.06 | 987.372 | 1000.43 | 1034.13 |
| **Protein (kCal)** | 5% | 7% | 10% | 20% | 45% |
| **Carbohydrate (kCal)** | 75% | 73% | 70% | 60% | 35% |
| **Fat (kCal)** | 20% | 20% | 20% | 20% | 20% |
| **Total (kCal/g)** | 4000 | 4000 | 4000 | 4000 | 4000 |
